# Exon Targeted Retrieval and Classification Toolbox (ExTRaCT): a gene search pipeline to find APOBEC3 Z-domains in novel bat genomes

**DOI:** 10.64898/2026.03.15.711917

**Authors:** Brenda Delamonica, Bat1K 21-Families Group, Mani Larijani, Thomas MacCarthy, Liliana M. Dávalos

## Abstract

**Motivation:** Several computation gene search tools exist to identify and annotate an ever-growing body of newly sequenced genomes of different species. Many annotation tools, however, fall short when the target species diverges from well-studied model organisms, and when searching for short genes with multiple copies.

**Results:** We have developed the Exon Targeted Retrieval and Classification Toolbox, ExTRaCT, an automated pipeline to identify any gene exon with conserved structure in novel species genome assemblies. In the use cases presented here, we applied our search tool to 102 bat genomes to find APOBEC3 gene family members. We show that our homolog search algorithm is efficient (run time average of 5 hours for over 100 genomes), works well with reference sequences distantly related to the target (1 out of 498 misclassifications, 0 false positives and 2 false negatives), and is easy to use. As genomic sequencing becomes faster and more accessible, ExTRaCT has downstream applications in phylogenetic, biochemical and genomic studies. It is a simple computational tool that provides a solution to target gene identification, requiring neither whole-genome-assembly annotations, nor prior knowledge of closely related species.

**Availability:** https://doi.org/10.5281/zenodo.15769018

**Supplementary information:** Supplementary data are available at *Bioinformatics* online.

## 1 Introduction

An initiative to sequence all living bat genomes, known as the Bat1K project, aims to discover unique adaptations in bats (Bat1K 21 Family Group, 2026; Morales et al.). However, existing computational tools to identify and annotate gene family members in non-model organism genome assemblies often rely on computationally expensive methods, substantive training, and fail to find small gene copies. Identifying gene family members is difficult in bats, which have evolved through numerous diversification events. Gene homology search tools, such as BLAST, rely on arbitrary hit thresholds which may lead to missing results, and subsequent manual analysis becomes tedious (Nestor et al., 2023). Machine learning annotation and gene prediction tools, such as TOGA (Tool to Infer Orthologs from Genome Alignments) (Kirilenko et al., 2023), can conduct full genome annotations, but may miss sections of complicated multi-domain sequences, especially when the target diverges from the reference, and training on new species requires computational expertise. Furthermore, full annotations may require isoform sequencing with long-read technology to identify complex or missing gene homologues based on RNA, which may not be available for purely computational studies (Jebb et al., 2020). In the case of small antimicrobial peptides in bats, MAKER (Cantarel et al., 2008), an integrated gene prediction pipeline, could only find single gene copies and omitted short multi-genes (Castellanos et al., 2023). To address gaps in pipelines that optimize for single-copy homologues, we introduce the Exon Targeted Retrieval and Classification Toolbox (ExTRaCT), a streamlined fully computational gene family search pipeline which integrates homologue search tools with a custom algorithm that can accurately identify sequences, relies on a small set of input reference data, requires little experience with complicated techniques and unburdens the user from excessive manual post evaluation (Delamonica, 2025).

The Apolipoprotein B mRNA Editing Catalytic Polypeptide-like (APOBEC) family of cytidine deaminating proteins plays an important role in mammalian immunity (Conticello, 2008; Harris & Dudley, 2015; Pecori et al., 2022). A3s are interferon-stimulated genes (ISG) that act on DNA or RNA viruses by causing C > T point mutations at hotspot motifs to cause changes to viral protein that may be deleterious (Pecori et al., 2022; Stavrou & Ross, 2015). Viruses with A3 mutational signatures, or A3 mutational signatures, or “footprints,” have been observed in several human exogenous virus such as herpesviruses, polyomaviruses, hepadnaviruses, and papillomaviruses (Poulain et al., 2020; Shapiro, Krug, & MacCarthy, 2021; Willems & Gillet, 2015) as well as endogenous retroviruses or retroelements (Anwar, Davenport, & Ebrahimi, 2013; Ito, Gifford, & Sato, 2020), zoonotic diseases that have spilled over to humans, including Mpox (Delamonica et al., 2023; Forni et al., 2023; O’Toole et al., 2023) and SARS-CoV-2 (Begum et al., 2024; Kim et al., 2022; Poulain et al., 2020). However, A3s are also known to be inhibited or antagonized by viral proteins (Cheng et al., 2019; Grant & Larijani, 2017; Li et al., 2023; Lochelt et al., 2005; Wood et al., 2009; Zhang et al., 2021). These findings have raised the question of whether A3 activity in the long term is proviral or antiviral, since A3 can play a role in driving viral evolution (Jonathan & Ikeda, 2023). As a result, the endemic pattern of human A3 mutation after viral spillover can *increase* viral diversity (Stavrou & Ross, 2015). A3s are therefore an important immune spillover barrier, albeit one with varying outcomes (Uriu et al., 2021). By understanding A3 characteristics in species known to harbor viruses, such as bats, we may better predict the mutational landscape of disease agents with zoonotic potential.

Bats are known hosts of viral agents that, when spilled over to other mammals, can produce zoonosis and pathogenicity (Guth et al., 2022; Irving et al., 2021). An extremely diverse mammalian order with over 1,500 living species that make up more than 20% of extant mammals spanning the globe, bats have varying physical and ecological phenotypes (Frick, Kingston, & Flanders, 2020; Martínez-Fonseca et al., 2024; Teeling et al., 2005). Efforts to understand bats as viral reservoirs and predict zoonosis come from previous zoonotic spillovers involving viruses that have relatives that circulate or are directly hosted in various bat species, such as Nipah, Marburg, Hendra, SARS and MERS (Becker et al., 2022; Becker et al., 2023; Forero-Munoz et al., 2024; Irving et al., 2021; Pandit et al., 2023; Streicker & Gilbert, 2020; Weber et al., 2023). Bat species vary in the expansion or contraction of immunity genes such as interferons (INFs) (which induce ISGs like A3) (Clayton et al., 2024; Streicker & Gilbert, 2020), antimicrobial peptides (Castellanos et al., 2023), and major histocompatibility complex (MHC) (Moreno Santillan et al., 2021) in ways that are lineage-specific, associated with viral interactions(Vazquez et al., 2025), and sometimes because of differences in diet or diet-related microbiome evolution changing their interactions with potential pathogens. Bat-centered research on viral zoonosis is complicated by bat diversity, varying longevity, data biases and lack of longitudinal data (Carlson et al., 2025; Streicker & Gilbert, 2020; Weber et al., 2023). Nevertheless, two groundbreaking studies (Hayward et al., 2018; Jebb et al., 2020) have endeavored to study A3s in bats, positing that increased copies of A3 in bats may contribute to virus resistance.

The first study (Hayward et al., 2018) analyzed the A3s of two bat species in the Pteropodidae family. They biochemically tested A3 gene expression, mutational activity and viral restriction from transcriptomes identified using BLAST. They found up to 13 possible genes in *Pteropus alecto*. For context, humans have 7 different A3s (A3A/B/C/D/F/G/H) and mice have a single copy (Salter, Bennett, & Smith, 2016). More recently, Jebb et al. (2020) examined six bat species, hereafter referred to as the “six bats” (*Rousettus aegyptiacus* in Pteropodidae and *Rhinolophus ferrumequinum* in Rhinolophidae, both in Yinpterochiroptera, *Phyllostomus discolor* in Phyllostomidae, *Molossus molossus* in Molossidae, and *Pipistrellus kuhlii* and *Myotis myotis* in Vespertilionidae, all in Yangochiroptera). The original six-bat study (Jebb et al., 2020) used isoform sequencing with long-read technology to supplement computational findings by TOGA to identify A3s. The species with the highest A3 count was *Pipistrellus kuhlii*, with a total of 8 distinct genes, although it is unclear if all are active or functional. In contrast, *Rousettus aegyptiacus* had 2 A3s, a marked difference in A3 count from *Pteropus*. It is therefore still unclear whether all bats have more A3s than other mammals or even more than primates, but as more bat genome assemblies become available, we need easy-to-use tools to extract A3 sequences to study their evolution and better understand their role in bat immunity.

Here we expanded these findings by searching for the catalytic Zinc coordinating domains (Z-domains) of A3s in 102 different bat species, including the six bats from Jebb et al. (2020), using ExTRaCT. Salter was the first to classify human cytidine deaminating Z-domains into four types, zinc-dependent deaminase sequence motif (ZDD), attributed to Activation induced deaminase (AID), and the APOBECs A1, A2 and A4, and the three A3 Z-domains, Z1, Z2 and Z3 (Salter, Bennett, & Smith, 2016). A3s can have either one or two catalytic Z-domain regions, while the rest have a single ZDD. Along with theories of expansion events in A3 Z-domains among bats, both Hayward and Jebb propose new bat-specific Z-domain structures, diverging from the canonical motifs described in Salter, Bennett and Smith (2016). Not only is ours the largest study of bat A3 Z-domains to date, but the bat assemblies analyzed include species from all 21 bat families which could further support evolutionary associations with immunity and introduce novel definitions of bat-specific A3 Z-domain motif structures.

Our results support the idea that A3 genes are expanded in some bat lineages. We used freely available computational tools and built ExTRaCT to streamline the process of finding genes in gene families in an easy-to-use suite of Python functions so that it may be used to find other structurally conserved genes for any species. Our algorithm is fast, does not require high computational power or expertise and has lenient user-defined search criteria to facilitate modifications and ensure accuracy of results. We show that our pipeline correctly found the desired sequences and identified copies previously missed by TOGA from the six-bat study. Considering the drawbacks of existing gene family annotation tools, our novel suite, ExTRaCT, has applications in evolutionary, phylogenetic or functional study of genes families beyond bat A3.

## 2 Methods

Assemblies from 103 genomes were downloaded from the Bat1K database (Bat1K 21 Family Group, 2026; Morales et al.). One species, *Myotis thysanodes*, was excluded from the analysis because the scaffolds were incomplete and none of the results passed quality control. Reference homologue A3 gene nucleotide FASTA files can be found in (NCBI Resource Coordinators, 2016) and the Hiller Lab databases (Jebb et al., 2020; Perez et al., 2025; UCSC, 2025). All data sources and accession numbers are in Supplementary Materials.

The ExTRaCT pipeline was written in Python version 3.7.10 (Python Software Foundation, 2021) and requires both Biopython (v1.81) modules SeqIO and Align IO (Cock et al., 2009), PyBedTools (v 0.10.0) (Dale, Pedersen, & Quinlan, 2011), HMMER version 3.1b2 (Eddy, 2011). EMBOSS (specifically the program getorf) (v6.6) (Rice, Longden, & Bleasby, 2000), Scipio (v1.4.1) (Keller et al., 2008)MAFFT (v 7.490) (Katoh & Standley, 2013), and RAxML (v8.2.18) (Stamatakis, 2014). The custom functions in ExTRaCT are prep_hmmer.py, which converts a given FASTA to the appropriate input HMMER profile structure for the main algorithm, Gene_search.py, and three post processing functions, Scipio_run.py, table_to_fasta.py and gene_Tree.py, which help analyze the results. ExTRaCT is efficient as it relies on limited information about the gene family and replaces manual activities with programmable automation.

Our approach begins by generating a reference homologue gene profile required for ExTRaCT. We used HMMER (v3.1b2) (Eddy, 2011) instead of BLAST (v2.9) (Camacho et al., 2009) to find homologous sequences because BLAST yielded fewer viable hits and required more manual steps to filter out target non-overlapping sequences. The core search algorithm (Gene_search.py) has four crucial steps which provide nucleotide and amino acid sequences that closely match previously known gene structures in a novel species where gene annotations may not yet be known or provided. The steps are: (1) find homologous sequence start and stop locations, (2) extract the sequence from the genome assembly, (3) identify open reading frames in the sequence and (4) filter protein sequences that match with a given structure. In the first step we use the HMMER function nhmmer to search for nucleotide homologues based on a profile created using hmmbuild (Eddy, 2011). Of these, the HMMER homologue search is the most time-consuming step in the process as it searches through the entire genome. The homologue search may also take longer if there are more known sequences in the FASTA homologue input file. HMMER returns start and end locations of homologues as a bed file, which need to then be extracted by PyBedTools (v0.8.1) (Dale, Pedersen, & Quinlan, 2011), the second step of our pipeline. Given a series of nucleotide sequences, ExTRaCT identifies which hits match a gene protein structure by identifying open reading frames (ORF) (EMBOSS v6.6 getorf) (Rice, Longden, & Bleasby, 2000) within a nucleotide sequence. However, true start and stop locations may change at step (3) and be different from the initial HMMER homologue search results. Lastly, sequences containing the canonical Z-domain motif are saved based on categorizations defined by the user. To ensure all gene copies are extracted, the result list includes all sequences, even duplicated non overlapping sequences within the same species. Overall, the main algorithm can take approximately 5 hours for over 100 species, depending on the size of the input files. However, once the ORF files are generated, there is an option to only test filtering of various motif structures, as ExTRaCT will automatically skip steps 1-3 if the HMMER, bed and ORF files already exist in the expected location. These first three steps do not depend on the motif category file that finally filters the gene hits. That way, a single test to try different categorization methods is easy, requires less memory and can take as little as 5 minutes for approximately 100 species. To optimize processing a large scale of species targets, it is best to split the input FASTA file with target genome paths and run smaller batches in parallel. Sample code is provided in the GitHub repository.

To test performance, we compared hit counts base on inputs by 1) reference species taxa (bat, primate and laurasiatherian), 2) domain specification (single, double or both) and 3) user-designed categorization based on Salter, Hayward and Jebb (motif sets A, B and C, in order). We report three different input strategies for a total of 27 tests in Table 1. We conducted a pairwise comparison for all 27 tests whose summaries can be found in Supplementary Materials. To test accuracy, we then compared the results to known bat A3s found in Jebb et al. (2020).

**Table 1.** Count of A3 Z-domains found in 102 bat genomes for 27 input tests.

| Input Clade | Domain Type | A (Salter) | B (Hayward) | C (Jebb) |
| --- | --- | --- | --- | --- |
| Bat | Single | 121 | 123 | 496 |
|  | Double | 120 | 122 | 489 |
|  | Both | 121 | 123 | 496 |
| Primate | Single | 120 | 122 | 495 |
|  | Double | 121 | 120 | 471 |
|  | Both | 121 | 123 | 495 |
| Laurasiatheria | Single | 120 | 122 | 487 |
|  | Double | 121 | 123 | 496 |
|  | Both | 121 | 123 | 496 |

ExTRaCT sequence counts of A3 Z-domain search conducted on 102 novel bat genome assemblies grouped by different input options: species homologue input, domain type and labeling protocol (A, B and C). “Both” means the input includes single and double domain A3s.

There are three post processing functions that finetune nucleotide sequence results using Scipio (v1.4.1) (Keller et al., 2008) (custom function Scipio_run.py), convert tables with amino acid and nucleotide sequences into FASTA files (custom function table_to_fasta.py), and generate multiple sequence alignments using MAFFT (v7.490) (Katoh & Standley, 2013) and infer phylogenetic using trees RAxML (v.8.2.18) (Stamatakis, 2014). We chose to keep the Scipio nucleotide track separate from the main algorithm since many pertinent analyses may be conducted with amino acid structure alone. However, we provide the Scipio code to extract accurate nucleotide sequence information, as the start and stop bounds are sometimes lost in step (3) of Gene_search.py and not updated when running getorf. The corrected nucleotide sequences could later be used to express and synthesize proteins to conduct experiments of gene activity and function. The optional post-processing step to examine phylogenetic trees of the output helps to validate which genes may be misclassified. These trees could be used for comparative phylogenetic studies that model the duplication and loss processes by which a given gene has evolved. Both Scipio and tree steps can be skipped if primary protein structure is all that is required.

Because of the known divergence between A3s among bats, we provide a final gene tree of amino acid sequences, rooted after phylogenetic inference using known AID genes as outgroups in FigTree (v1.4.4) (Rambaut, 2018). False positive, or non-homologues, results can be identified if sequences form distinct clades that render a known gene subfamily paraphyletic or polyphyletic (e.g., by being nested within the outgroup clade). In contrast, a false-negative results would be a gene omitted from the results of a known species, which we evaluated manually with the six bats (Jebb et al., 2020). However, misclassification errors can be found using a species tree if a clade comprises different gene categories, i.e., results yield non monophyletic groups. While this analysis can identify homologues, we are unable, without proper phylogenetic comparison to the species tree, to distinguish paralogs (which derive from gene duplication) or orthologs (which derive from speciation). In Figure 1 we removed the AID outgroup and kept all the APOBECs, (A1, A2 and A4 in grey). To test if any of the sequences have been mislabeled and belong to different domain groups, we verify that each of the domains form clades as well.

**Fig. 1.**
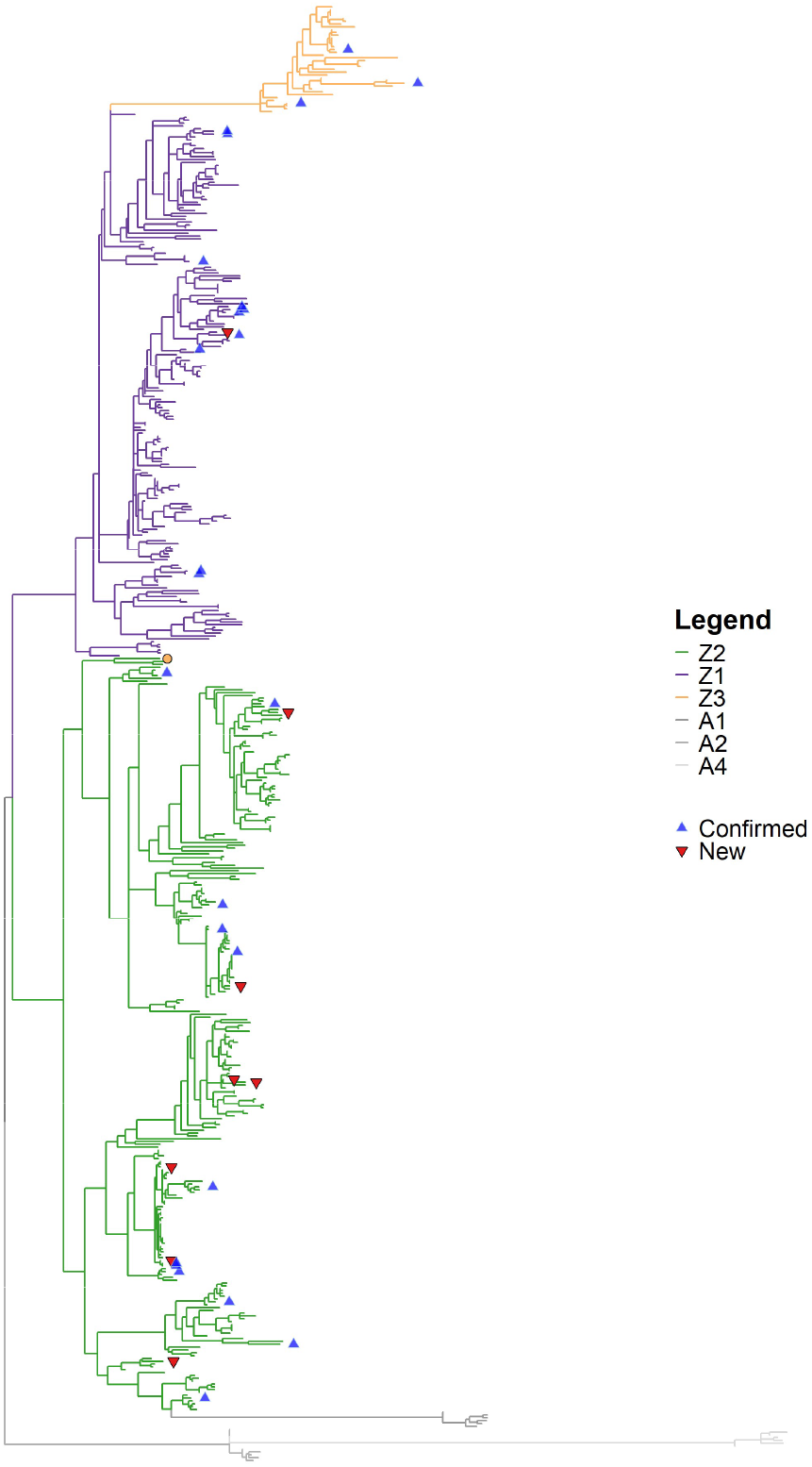
Gene tree of bat A3 Z-domains. Phylogenetic tree with the final 498 sequences from ExTRaCT pulled from 102 otherwise unannotated bat genomes. Outgroup of known six bat ZDDs of the APOBECs A1, A2 and A3 in grey. Different Z-domain subfamily members are labeled by color (Z1 is purple, Z2 is green and Z3 is orange). Red upside-down triangles indicate hits which are not previously annotated Z domains of known A3s. Blue triangles indicate confirmed A3 Z domains of known six bats.

Various packages in R (v4.3.2) (R Core Team, 2023) were used to plot the phylogenetic tree (gtree (v3.10.0) (Yu, 2020; Yu et al., 2020), ggplot2(v3.5.1) (Wickham et al., 2024), phytools(v2.1-1) (Revell, 2024), ape (v5.7-1) (Paradis & Schliep, 2019), ggmsa (v 1.8.0) (Zhou et al., 2022), Biostrings (2.70.1) (Pagès et al., 2023), seqinr (v4.2-36) (Charif & Lobry, 2007) and msa (v1.34.0) (Bodenhofer et al., 2015)). All tables were reformatted in R using the packages dplyr (v1.1.4) (Wickham et al., 2023), tidyr (v1.3.0) (Wickham, Vaughan, & Girlich, 2023), openxlsx (v4.2.7.1) (Schauberger & Walker, 2024), readxl (v1.4.3) (Wickham & Bryan, 2023), writeexl (v1.5.1) (Ooms, 2024) and purrr (v1.0.2) (Wickham & Henry, 2023).

## 3 Results

We tested three input strategies, compared the best results to known bat annotations and inferred a gene tree for all 102 bat A3Z genes identified using ExTRaCT.

### 3.1 ExTRaCT is robust across different mammalian clades

When considering different clade inputs, for the same domain type and motif set, closely related species should result in similar hits. For domain type both the counts across the different clades are equal under motif set A (121) and B (123) but for motif set C the primate test is missing one hit (primate = 495, chiroptera and laurasiatherian = 496). ExTract has a very low error rate (1/496) when using a different input clade (primate) than the target species, it is otherwise robust. The input sequences need not be of the same order as the target species to garner the same results. Input sequences of the same superorder are only slightly preferred. This is beneficial if no known relatives to a target species have gene information or annotated genomes.

While the counts may be the same, the actual sequences found might differ, so we refer to a pairwise match inspection for all test scenarios (Supplementary Materials). We verified that Laurasiatheria and bat domain type both and motif set C input tests contained all the sequence hits, which we elaborate on in the next section. Furthermore, testing multiple domain types is A3-specific. Catalytic domain configurations vary greatly across species and to capture the correct sequences we found that including all genes (both) is best. For the rest of the analysis and for brevity, we focus on domain type ‘both.’

### 3.2 Motif definition is the most critical and flexible input

What impacts the final count the most is the last step of Gene_search.py which filters and categorizes hits that match a user-provided labeling criterion, motif sets A, B and C. In our test, motif set A has the fewest hits. This may be because the canonical motifs were based on human A3s, have fewer domain categories than those that have since been seen in bats, and are more restrictive protein structures than in Hayward and Jebb. Despite this, motif set B only captured two more sequences than set A. It is possible that despite the inclusion of a new Z-domain in motif B, other changes are more restrictive, limiting the final output. Also, since motif B is only based on two bat species in the same family, and given the known diversity among bats, these categories may not capture sequence diversity of the whole order. The difference in results between motif set C and either motif set A or B is remarkable. Across all test scenarios anywhere from 373-375 more sequences were found in the motif C set test result list.

We found in our pairwise match inspection that two sequences in *Furipterus horrens* were found in the Laurasiatheria reference motif set B test but neither in Laurasiatheria nor in bat motif set C. We verified that this is due to the differences in structure of the Z2B domain motif between Hayward and Jebb. Out of all 27 pairwise inspections, these two sequences were the only sequences that are missing completely from the combined domain Laurasiatheria or bat motif set C test. We consider these false negative results for this test and add them manually to produce our final list of 498 genes across the 103 bat species (Supplementary Materials).

### 3.3 Validating results with known bat A3s

To ensure that our algorithm finds true A3 Z-domains, and identify false negatives, we count the number of known Z-domains from the six bat species in Jebb et al. (2020) that were found using ExTRaCT in **Error! Reference source not found**.. Motif set C tests returned all the 25 known Z-domains across all clade and domain type tests. However, motif set A and B tests had the same results, with a high rate of false negatives (22/25) and only 3 matching Z-domains to known bat A3s. This further emphasizes the importance of motif categorization. Motif C returns no false negatives because it is the most up to date and relevant structure for bats in different families.

In the final gene list, we note that among the six bat species, an additional eight sequences were found, different from the 25 known bat A3 Z-domains found using TOGA. These may be A3 paralogs or orthologs, depending on when and how the genes diverged. The sequences may be copies from duplication events or speciation events, and they may be non-coding sequences or nonfunctional genes. This is a primary example of the benefit of ExTRaCT over other annotation techniques to better understand the evolution of gene families, as it can find gene family members other conventional computational annotation methods miss.

### 3.4 Producing a reliable gene list for phylogenetic evolutionary analysis and novel Z-domain discovery in bats

A typical step when conducting a manual gene homology search is to align the sequences, infer a phylogeny and validate hits are in the correct gene family or subfamily. We built table_to_fasta.py and gene_Tree.py to automate this post-processing. This results in Figure 1, a phylogeny of the 498 ExTRaCT sequence hits from the Laurasiatheria combined domain motif set C test with two added sequences (false negative results). Known bat cytidine deaminases are the outgroup (A1, A2, A4=grey). The different Z-domain categories are labeled by color (Z1=purple, Z2=green, Z3=orange). None of the sequences form a clade with the outgroup so we can be confident that all results are monophyletic. This is probably because our algorithm focuses on filtering out by structure rather than a heuristic similarity score threshold, enforcing finer structure similarity.

All the sequences, except one tip labeled Z3 (orange dot) in the Z2 clade, form a clade with the correctly labeled domain categories. This sequence is in the *Nycteris thebaica* (Nt1) assembly and is the second Z3 domain found in that species. Having two Z3 domains is rare (Munk, Willemsen, & Bravo, 2012; OhAinle et al., 2008) as they are thought to be genotoxic (Ito, Gifford, & Sato, 2020; Suspene et al., 2011) so it is likely that this sequence is mislabeled and a possible misclassification error. In an MSA of the Z3 domains (Supplementary Materials), Nt1 has a unique structure, unlike that of other Z3s, and is unlike any of the Z-domain sequences identified in bats so far. It has both a TWS triplet, a defining distinction of Z3 domains, and a WF couplet in position 9-10, which is also an indicator of Z2. It is possible this is either a result of recombination or some other evolutionary event. Since most of the sequence is clearly not well aligned to other Z3s and aligns more closely to other Z2s, we relabel it to Z2 in our final gene list. However, this raises the possibility that bats have even more distinct Z-domain types than defined most recently by Jebb et al. (2020).

The 25 known six bat species Z-domain sequences are labeled in brown triangles and the 8 possible paralog or orthologs are labeled by empty pink triangles. We note that many of the known genes and ExTRaCT-identified homologs are closely related, but without a gene tree-species tree reconciliation it is not clear which are paralogs, and which are orthologs. A gene tree-species tree reconciliation could identify duplication, speciation and loss events across lineages over evolutionary time. For example, from our gene list we see that count of Z-domains varies drastically by species, where total counts range from 1-23 total Z-domain hits per species. A phylogenetic reconciliation analysis would shed some light into the evolutionary relationships between the species and A3 genes. Further work could relate these evolutionary dynamics to immunity and host-virus associations.

### 3.5 ExTRaCT is fast and scalable

Our core algorithm in the pipeline is fast and does not require much memory. Depending on the size of the inputs, a single genome assembly can be analyzed in 2-3 minutes. The largest genome assembly in our test case was *Saccopteryx bilineata*, approximately 2.5 Gb. For a single run, HMMER is the longest step, and it takes around 2 minutes. The subfunctions, pybedtools, getorf and our filtering code, all took less than a minute. The total time to run our pipeline on a genome of 2.5Gb in size was 3 minutes. Depending on computing capabilities, at scale, the run of 103 genomes took an average of 5 hours. Genome sequencing is ubiquitous and there is a growing stream of novel genomes becoming publicly available every day. Out of 498 sequences, our algorithm only misclassified one sequence, demonstrating a low misclassification error rate, zero false positives (each gene subfamily is monophyletic) and two false negatives (found in other tests). Having fast, accurate and easy-to-use methods to conduct targeted annotations of gene families is very beneficial as the landscape of data availability increases.

This order-of-magnitude scaling in annotation of bat A3s has not been done before, and we demonstrate that as genome assemblies become more available, ExTRaCT may be used at a massive scale. There are more genomic data than ever before, but to garner insights into the evolution of gene families requires methods to identify genes. We chose to investigate A3s, a gene family at the forefront of immunity and virus evolution, in bats, a highly diverse clade that includes several species known to be reservoirs of viruses with zoonotic potential and in which A3s appear to have undergone highly dynamic evolutionary changes. With ExTRaCT, others can start to identify A3s or any other salient genes, study the evolution of a gene family or characterize gene expression and classify gene activity. In our case molecular biologists can use our bat A3 results to study A3 and virus coevolution to continue to answer the question of the role A3 mutagenesis plays, whether anti- or pro-viral, in bats. These types of studies could improve the prediction of mutagenesis and recognize viruses with potential risk to humans. By identifying the catalytic regions of bat A3 genes, our case study is the first step in understanding how A3s are involved in the exceptional immunity of bats.

