## Supplementary Materials for "Exon Targeted Retrieval and Classification Toolbox (ExTRaCT): a gene search pipeline to find APOBEC3 Z-domains in novel bat genomes"

### Supplementary Materials (SM)

#### SM1 Background on Bat Immunity and Small Gene Annotation

Some believe that bats have heightened immunity because the sustained temperature held during flight contributes to positive selection on DNA damage and repair genes linked to enhanced immunity and disease tolerance (Banerjee et al., 2020; O'Shea et al., 2014; Zhang et al., 2013). While this hypothesis is often cited as an explanation for certain increases by gene duplication in bat immunity genes, it does not account for other demographic or behavioral differences that can shape the immune system such as roosting, diet, geographical realm or exposure to disease. Anthropogenic stressors such as habitat loss and food scarcity amplify bat originating spillover, posing a zoonotic risk to humans (Becker et al., 2023; Carlson et al., 2025; Eby et al., 2023). Our work is motivated by the question of whether variation in A3 Z-domain counts impact bat immunity.

The first step to answer this question through phylogenetic, structural and functional analysis requires robust tools to accurately identify gene families in a target genome assembly. Existing automated methods for genome annotation may overlook or misrepresent gene family member targets, may rely on advanced or computationally burdensome techniques and may be limited to reference genomes of fully annotated model organisms. We mention in the man text examples with TOGA and MAKER, but another annotation tool AUGUSTUS (Stanke & Morgenstern, 2005) which has been trained on many species, requires computational expertise. Additionally, TOGA raw output (Perez et al., 2025) can return missing or partial sequences, requiring noncomputational protocols to make up for this deficit. Together or in some combination, these shortcomings can mislead characterization of gene families. To address these limitations, biologists often have their own manual processes to validate gene searches, with varying degrees of automation. Here we provide an easy-to-use pipeline that is fully computational, filling gaps in traditional out-of-the box tools to enable downstream gene family studies.

#### SM2 Input sequence and genome data

##### SM2.1. Table. Accession numbers, source and date downloaded

Names, accession numbers or other identifiers and date of download of (1) reference sequence APOBEC3 (from NCBI (NCBI Resource Coordinators, 2016)), (2) target species genomes (from globus (Bat1K 21 Family Group, 2026)) and (3) six bat A3 (from UCSC senckenberg genome browser (UCSC, 2025))

| Species | Sequence Name | Source | #Accession or file name | Date downloaded |
|---|---|---|---|---|
| *Macaca mulatta* | A3C | NCBI | EU381233.1 | 11/18/2024 |
| *Chlorocebus aethiops* | A3C | NCBI | EU381232.1 | 11/18/2024 |
| *Homo sapiens* | A3A (v1) Z1 | NCBI | NM_145699.4 | 11/18/2024 |
| *Homo sapiens* | A3B (v1) Z2-Z1 | NCBI | NM_004900.5 | 11/18/2024 |
| *Homo sapiens* | A3C Z2 | NCBI | NM_014508.3 | 11/18/2024 |
| *Homo sapiens* | A3D (v1) Z2-Z2 | NCBI | NM_152426.4 | 11/18/2024 |
| *Homo sapiens* | A3F (v1) Z2 – Z2 | NCBI | NM_145298.6 | 11/18/2024 |
| *Homo sapiens* | A3F (cds) Z2 – Z2 | NCBI | JF262037.1 | 11/18/2024 |
| *Homo sapiens* | A3G (cds) Z2-Z1 | NCBI | JF262036.1 | 11/18/2024 |
| *Homo sapiens* | A3H (vSV-154) Z3 | NCBI | NM_001166004.3 | 11/18/2024 |
| *Bos taurus* | A3 Z1 | NCBI | EU864534.1 | 11/18/2024 |
| *Bos taurus* | A3Z2 | NCBI | EU864535.1 | 11/18/2024 |
| *Bos taurus* | A3Z3 | NCBI | EU864536.1 | 11/18/2024 |
| *Equus caballus* | A3 Z2a-Z2b | NCBI | FJ527822.1 | 11/18/2024 |
| *Equus caballus* | A3 Z2c-Z2d | NCBI | FJ527823.1 | 11/18/2024 |
| *Equus caballus* | A3 Z3 v1 | NCBI | FJ532286.1 | 11/18/2024 |
| *Equus caballus* | A3 Z1b | NCBI | FJ532287.1 | 11/18/2024 |
| *Equus caballus* | A3 Z2e | NCBI | FJ532288.1 | 11/18/2024 |
| *Equus caballus* | A3 Z3 | NCBI | FJ532289.1 | 11/18/2024 |
| *Felis catus* | A3 Cc | NCBI | EU109281.1_cds_ABW83271.1_1 | 11/18/2024 |
| *Felis catus* | A3 Ca | NCBI | EU109281.1_cds_ABW83272.1_2 | 11/18/2024 |
| *Felis catus* | A3 Cb | NCBI | EU109281.1_cds_ABW83273.1_3 | 11/18/2024 |
| *Felis catus* | A3H | NCBI | EU109281.1_cds_ABW83274.1_4 | 11/18/2024 |
| *Felis catus* | A3 | NCBI | GU097660.1 | 11/18/2024 |
| *Lynx lynx* | A3 | NCBI | GU097661.1 | 11/18/2024 |
| *Ovis aries* | A3 Z1 | NCBI | EU864541.1 | 11/18/2024 |
| *Ovis aries* | A3 Z2 | NCBI | EU864542.1 | 11/18/2024 |
| *Ovis aries* | A3 Z3 | NCBI | EU864543.1 | 11/18/2024 |
| *Panthera leo bleyenberghi* | A3 | NCBI | GU097662.1 | 11/18/2024 |
| *Panthera tigris corbetti* | A3 | NCBI | GU097663.1 | 11/18/2024 |
| *Puma concolor* | A3 | NCBI | GU097659.1 | 11/18/2024 |
| *Sus scrofa* | A3 Z2 | NCBI | EU864539.1 | 11/18/2024 |
| *Sus scrofa* | A3 Z3 | NCBI | EU864540.1 | 11/18/2024 |
| *Sus scrofa* | A3 Z2-Z3 | NCBI | EU871586.1 | 11/18/2024 |
| *Sus scrofa* | A3 Z2_1 | NCBI | FJ716801.2 | 11/18/2024 |
| *Molossus molossus* | A3A | UCSC | HLmolMol2 APOBEC3A | 10/17/2024 |
| *Molossus molossus* | A3A2 | UCSC | HLmolMol2 APOBEC3A2 | 10/17/2024 |
| *Molossus molossus* | A3C | UCSC | HLmolMol2 APOBEC3C | 10/17/2024 |
| *Molossus molossus* | A3H | UCSC | HLmolMol2 APOBEC3H | 10/17/2024 |
| *Myotis myotis* | A3A | UCSC | HLmyoMyo6 APOBEC3A | 10/17/2024 |

| | | | | |
|---|---|---|---|---|
| *Myotis myotis* | A3A2 | UCSC | HLmyoMyo6 APOBEC3A2 | 10/17/2024 |
| *Myotis myotis* | A3A3 | UCSC | HLmyoMyo6 APOBEC3A3 | 10/17/2024 |
| *Myotis myotis* | A3C | UCSC | HLmyoMyo6 APOBEC3C2 | 10/17/2024 |
| *Myotis myotis* | A3C2 | UCSC | HLmyoMyo6 APOBEC3C2 | 10/17/2024 |
| *Pipistrellus kuhlii* | A3A | UCSC | HLpipKuh2 APOBEC3A | 10/17/2024 |
| *Pipistrellus kuhlii* | A3A2 | UCSC | HLpipKuh2 APOBEC3A2 | 10/17/2024 |
| *Pipistrellus kuhlii* | A3A3 | UCSC | HLpipKuh2 APOBEC3A3 | 10/17/2024 |
| *Pipistrellus kuhlii* | A3A4 | UCSC | HLpipKuh2 APOBEC3A4 | 10/17/2024 |
| *Pipistrellus kuhlii* | A3A6 | UCSC | HLpipKuh2 APOBEC3A6 | 10/17/2024 |
| *Pipistrellus kuhlii* | A3C | UCSC | HLpipKuh2 APOBEC3C | 10/17/2024 |
| *Pipistrellus kuhlii* | A3C2 | UCSC | HLpipKuh2 APOBEC3C2 | 10/17/2024 |
| *Pipistrellus kuhlii* | A3C3 | UCSC | HLpipKuh2 APOBEC3C3 | 10/17/2024 |
| *Phyllostomus discolor* | A3A | UCSC | HLphyDis3 APOBEC3A | 10/17/2024 |
| *Phyllostomus discolor* | A3A2 | UCSC | HLphyDis3 APOBEC3A2 | 10/17/2024 |
| *Phyllostomus discolor* | A3C2 | UCSC | HLphyDis3 APOBEC3C2 | 10/17/2024 |
| *Rousettus aegyptiacus* | A3 | UCSC | HLrouAeg4 APOBEC3A | 10/17/2024 |
| *Rousettus aegyptiacus* | A3A | UCSC | HLrouAeg4 APOBEC3A | 10/17/2024 |
| *Rhinolophus ferrumequinum* | A3A | UCSC | HLrhiFer5 APOBEC3A | 10/17/2024 |
| *Rhinolophus ferrumequinum* | A3A2 | UCSC | HLrhiFer5 APOBEC3A2 | 10/17/2024 |
| *Rhinolophus ferrumequinum* | A3B | UCSC | HLrhiFer5 APOBEC3B | 10/17/2024 |
| *Rhinolophus ferrumequinum* | A3C | UCSC | HLrhiFer5 APOBEC3A | 10/17/2024 |
| *Cistugo seabrae* | genome | Bat 1k | mCisSea1.1.hap1 | 10/18/2024 |
| *Craseonycteris thonglongyai* | genome | Bat 1k | mCraTho2.1.hap1 | 10/19/2024 |
| *Rhynchonycteris naso* | genome | Bat 1k | mRhyNas1.2.hap1 | 10/18/2024 |
| *Saccopteryx bilineata* | genome | Bat 1k | mSacBil1.1.pri | 10/18/2024 |
| *Saccopteryx leptura* | genome | Bat 1k | mSacLep1.1.pri | 10/18/2024 |
| *Taphozous melanopogon* | genome | Bat 1k | mTapMel1.1.pri | 10/18/2024 |
| *Furipterus horrens* | genome | Bat 1k | mFurHor1.1.pri | 10/18/2024 |
| *Aselliscus stoliczkanus* | genome | Bat 1k | mAseSto1.2.pri | 10/19/2024 |
| *Doryrhina cyclops* | genome | Bat 1k | mDorCyc1.2.pri | 10/19/2024 |
| *Hipposideros abae* | genome | Bat 1k | mHipAba1.1.hap1 | 10/19/2024 |
| *Hipposideros armiger* | genome | Bat 1k | mHipArm2.1.pri | 10/19/2024 |
| *Hipposideros caffer* | genome | Bat 1k | mHipCaf2.1.hap1 | 10/19/2024 |
| *Hipposideros jonesi* | genome | Bat 1k | mHipJon1.1.hap1 | 10/19/2024 |
| *Hipposideros larvatus* | genome | Bat 1k | mHipLar1.2.pri | 10/19/2024 |
| *Hipposideros swinhoii* | genome | Bat 1k | mHipSwi1.1.pri | 10/19/2024 |
| *Megaderma spasma* | genome | Bat 1k | mMegSpa1.1.pri | 10/19/2024 |
| *Miniopterus australis* | genome | Bat 1k | mMinAus1.1.pri | 10/18/2024 |
| *Miniopterus natalensis* | genome | Bat 1k | mMinNat1.1.hap2 | 10/18/2024 |
| *Miniopterus schreibersii* | genome | Bat 1k | mMinSch1.1.hap1 | 10/18/2024 |
| *Eumops nanus* | genome | Bat 1k | mEumNan1.1.hap1 | 10/18/2024 |
| *Molossus alvarezi* | genome | Bat 1k | mMolAlv1.1.hap1 | 10/18/2024 |
| *Molossus molossus* | genome | Bat 1k | mMolMol1.2.pri | 10/18/2024 |
| *Molossus nigricans* | genome | Bat 1k | mMolNig1.2.hap1 | 10/18/2024 |
| *Mops condylurus* | genome | Bat 1k | mMopCon1.1 | 10/19/2024 |
| *Tadarida brasiliensis* | genome | Bat 1k | mTadBra1.3.pri | 10/19/2024 |
| *Mormoops megalophylla* | genome | Bat 1k | mMorMeg1.1.pri | 10/18/2024 |
| *Mystacina tuberculata* | genome | Bat 1k | mMysTub1.2.pri | 10/18/2024 |
| *Myzopoda aurita* | genome | Bat 1k | mMyzAur1.1.pri | 10/18/2024 |
| *Natalus tumidirostris* | genome | Bat 1k | mNatTum1.1.pri | 10/19/2024 |
| *Noctilio leporinus* | genome | Bat 1k | mNocLep2.1.pri | 10/18/2024 |
| *Nycteris thebaica* | genome | Bat 1k | mNycThe1.1.hap1 | 10/18/2024 |
| *Artibeus intermedius* | genome | Bat 1k | mArtIn1.1.hap1 | 10/18/2024 |
| *Artibeus lituratus* | genome | Bat 1k | mArtLit1.1.hap1 | 10/18/2024 |
| *Brachyphylla cavernarum* | genome | Bat 1k | mBraCav1.1.hap1 | 10/18/2024 |
| *Carollia perspicillata* | genome | Bat 1k | mCarPer1.2.pri | 10/18/2024 |
| *Centurio senex* | genome | Bat 1k | mCenSen1.1.hap1 | 10/18/2024 |
| *Choeroniscus minor* | genome | Bat 1k | mChoMin1.1.hap1 | 10/18/2024 |
| *Desmodus rotundus* | genome | Bat 1k | mDesRot1.15.hap1 | 10/18/2024 |
| *Diaemus youngii* | genome | Bat 1k | mDiaYou1.3.hap1 | 10/18/2024 |
| *Diphylla ecaudata* | genome | Bat 1k | mDipEca1.2.hap1 | 10/18/2024 |
| *Ectophylla alba* | genome | Bat 1k | mEctAlb1.1.hap1 | 10/18/2024 |
| *Erophylla bombifrons* | genome | Bat 1k | mEroBom1.1.hap1 | 10/18/2024 |
| *Glossophaga mutica* | genome | Bat 1k | mGloMut1.1.hap1 | 10/18/2024 |
| *Glossophaga soricina* | genome | Bat 1k | mGloSor1.2.pri | 10/18/2024 |
| *Glyphonycteris daviesi* | genome | Bat 1k | mGlyDav1.1.hap1 | 10/18/2024 |

| | | | | |
|---|---|---|---|---|
| *Leptonycteris yerbabuenae* | genome | Bat 1k | mLepYer2.1.hap1 | 10/18/2024 |
| *Lionycteris spurrelli* | genome | Bat 1k | mLioSpu1.1.hap1 | 10/18/2024 |
| *Lonchorhina inusitata* | genome | Bat 1k | mLonInu1.1.hap1 | 10/18/2024 |
| *Macrophyllum macrophyllum* | genome | Bat 1k | mMacMac1.1.hap1 | 10/18/2024 |
| *Macrotus waterhousii* | genome | Bat 1k | mMacWat1.1.hap1 | 10/18/2024 |
| *Micronycteris megalotis* | genome | Bat 1k | mMicMeg1.1.pri | 10/18/2024 |
| *Phyllostomus discolor* | genome | Bat 1k | mPhyDis1.3.pri | 10/18/2024 |
| *Phyllostomus hastatus* | genome | Bat 1k | mPhyHas1.1.pri | 10/18/2024 |
| *Platyrrhinus guianensis* | genome | Bat 1k | mPlaGui1.1.hap1 | 10/18/2024 |
| *Rhinophylla pumilio* | genome | Bat 1k | mRhiPum1.1.hap1 | 10/18/2024 |
| *Trachops cirrhosus* | genome | Bat 1k | mTraCir3.1.hap1 | 10/18/2024 |
| *Trinycteris nicefori* | genome | Bat 1k | mTriNic1.1.hap1 | 10/18/2024 |
| *Uroderma convexum* | genome | Bat 1k | mUroCon1.1.pri | 10/18/2024 |
| *Vampyressa thyone* | genome | Bat 1k | mVamThy1.1.hap1 | 10/18/2024 |
| *Cynopterus sphinx* | genome | Bat 1k | mCynSph1.1.pri | 10/19/2024 |
| *Eonycteris spelaea* | genome | Bat 1k | mEonSpe1.2.hap1 | 10/19/2024 |
| *Hypsignathus monstrosus* | genome | Bat 1k | mHypMon1.1.pri | 10/19/2024 |
| *Rousettus aegyptiacus* | genome | Bat 1k | mRouAeg1.4.pri | 10/19/2024 |
| *Rhinolophus affinis* | genome | Bat 1k | mRhiAff1.2.pri | 10/19/2024 |
| *Rhinolophus ferrumequinum* | genome | Bat 1k | mRhiFer1.5.pri | 10/19/2024 |
| *Rhinolophus foetidus* | genome | Bat 1k | mRhiFoe1.2.pri | 10/19/2024 |
| *Rhinolophus hipposideros* | genome | Bat 1k | mRhiHip1.1.hap1 | 10/19/2024 |
| *Rhinolophus pearsonii* | genome | Bat 1k | mRhiPea1.1.pri | 10/19/2024 |
| *Rhinolophus perniger lanosus* | genome | Bat 1k | mRhiPer1.2.pri | 10/19/2024 |
| *Rhinolophus sinicus* | genome | Bat 1k | mRhiSin3.1.pri | 10/19/2024 |
| *Rhinolophus trifoliatus* | genome | Bat 1k | mRhiTri1.2.pri | 10/19/2024 |
| *Rhinolophus yonghoiseni* | genome | Bat 1k | mRhiYon1.2.pri | 11/12/2024 |
| *Triaenops persicus* | genome | Bat 1k | mTriPer1.1.hap1 | 11/12/2024 |
| *Rhinopoma microphyllum* | genome | Bat 1k | mRhiMic1.1.pri | 11/5/2024 |
| *Rhinopoma muscatellum* | genome | Bat 1k | mRhiMus1.1.pri | 11/12/2024 |
| *Thyroptera tricolor* | genome | Bat 1k | mThyTri1.1.pri | 10/18/2024 |
| *Antrozous pallidus* | genome | Bat 1k | mAntPal2.1.pri | 10/19/2024 |
| *Corynorhinus mexicanus* | genome | Bat 1k | mCorMex1.1.pri | 10/19/2024 |
| *Corynorhinus townsendii* | genome | Bat 1k | mCorTow1.1.hap1 | 10/19/2024 |
| *Eptesicus fuscus* | genome | Bat 1k | mEptFus1.3.pri | 10/19/2024 |
| *Eptesicus nilssonii* | genome | Bat 1k | mEptNil2.1.pri | 10/19/2024 |
| *Lasiurus ega* | genome | Bat 1k | mLasEga1.1.hap1 | 10/19/2024 |
| *Myotis auriculus* | genome | Bat 1k | mMyoAur1.1.pri | 10/19/2024 |
| *Myotis californicus* | genome | Bat 1k | mMyoCal1.1.pri | 10/19/2024 |
| *Myotis daubentonii* | genome | Bat 1k | mMyoDau2.1.pri | 10/19/2024 |
| *Myotis evotis* | genome | Bat 1k | mMyoEvo1.1.pri | 10/19/2024 |
| *Myotis lucifugus* | genome | Bat 1k | mMyoLuc2.1.pri | 10/19/2024 |
| *Myotis myotis* | genome | Bat 1k | mMyoMyo1.6.pri | 10/19/2024 |
| *Myotis mystacinus* | genome | Bat 1k | mMyoMys1.1.hap1 | 10/19/2024 |
| *Myotis nigricans* | genome | Bat 1k | mMyoNig1.1.pri | 10/19/2024 |
| *Myotis occultus* | genome | Bat 1k | mMyoOcc1.1.pri | 10/19/2024 |
| *Myotis pilosus* | genome | Bat 1k | mMyoPil1.1.pri | 10/19/2024 |
| *Myotis thysanodes* | genome | Bat 1k | mMyoThy1.1.pri | 10/19/2024 |
| *Myotis velifer* | genome | Bat 1k | mMyoVel1.1.pri | 10/19/2024 |
| *Myotis vivesi* | genome | Bat 1k | mMyoViv1.2.pri | 10/19/2024 |
| *Myotis volans* | genome | Bat 1k | mMyoVol1.1.pri | 10/19/2024 |
| *Myotis yumanensis* | genome | Bat 1k | mMyoYum1.1.hap1 | 10/19/2024 |
| *Nyctalus aviator* | genome | Bat 1k | mNycAvi1.1.pri | 10/19/2024 |
| *Pipistrellus kuhlii* | genome | Bat 1k | mPipKuh1.2.pri | 10/19/2024 |
| *Pipistrellus nathusii* | genome | Bat 1k | mPipNat1.1.hap1 | 10/19/2024 |
| *Pipistrellus pygmaeus* | genome | Bat 1k | mPipPyg1.1.pri | 10/19/2024 |
| *Plecotus auritus* | genome | Bat 1k | mPleAur1.1.pri | 10/19/2024 |
| *Vespertilio murinus* | genome | Bat 1k | mVesMur1.1.pri | 10/19/2024 |

#### SM3 Architecture Map

##### SM3.1. Figure. ExTRaCT small gene search pipeline

Pipeline architecture includes search algorithm and post processing steps. Each track corresponds to a custom Python command. The green rounded outline indicates an input file, the green diagonal and rounded outline indicates an output file that may be used as an input for the following track, and the blue squares indicates a process in the code. Preprocessing depends on hmmer and converts a FASTA file into a HMMER profile which is a required input for the gene search algorithm. The core gene search algorithm consists of four major processes with three program dependencies (hmmer, bedtools and getorf) and 3 required input files. The output is a table which can be processed into final FASTA and phylogenetic tree for analysis in the postprocessing tracks.

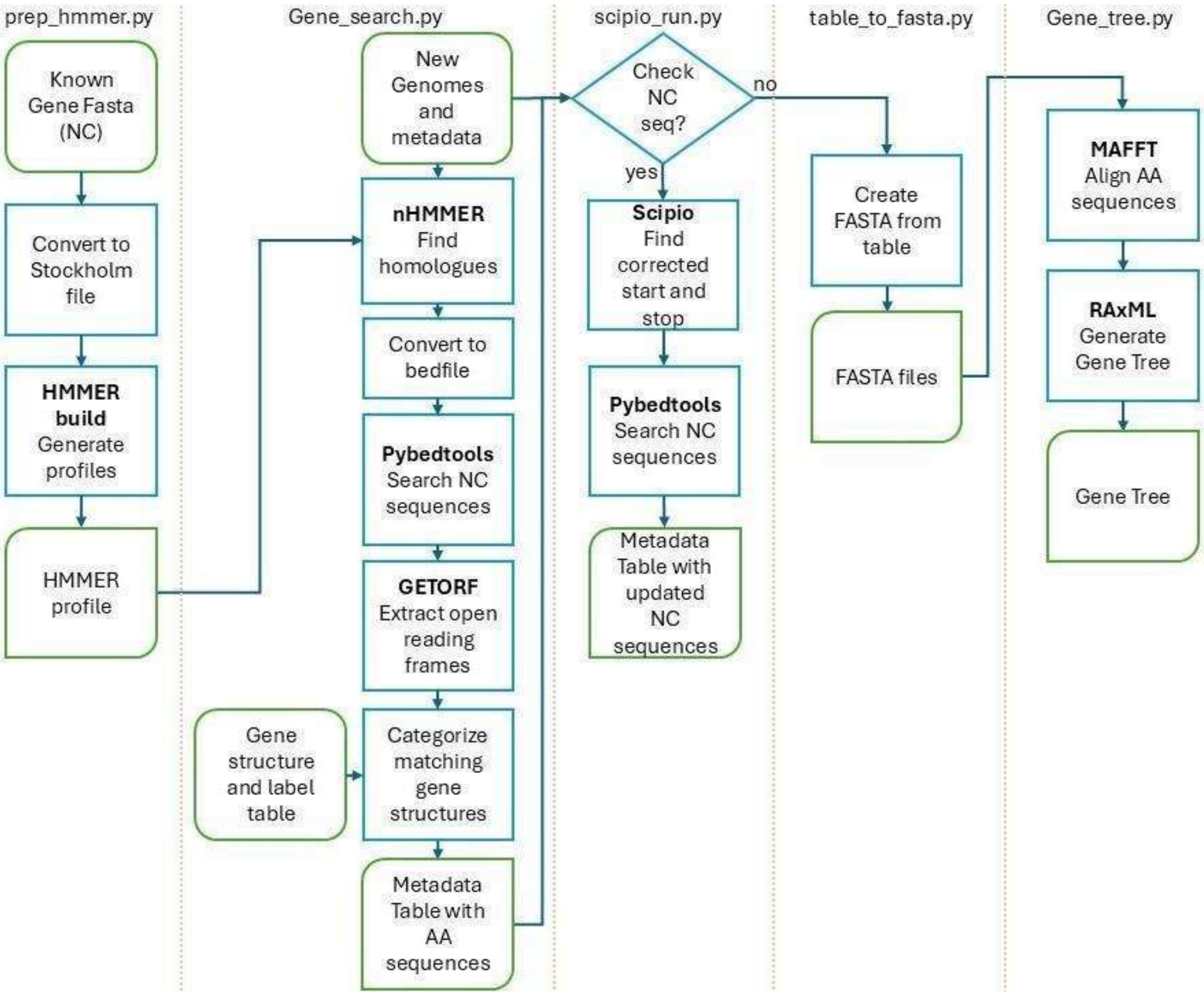


##### SM3.2. Program requirement details

Python version 3.7.10 (Foundation, 2021) (Anaconda is recommended but not required) Biopython (v1.81) SeqIO and Align IO (Cock et al., 2009) (pandas v.1.3.5 (Reback et al., 2021), regex/re 2024.4.16 (Hoyt, 2024), gzip, sys, os, argparse (Foundation, 2021)), PyBedTools (v 0.10.0) (Dale, Pedersen, & Quinlan, 2011), HMMER version 3.1b2 is required (Eddy, 2011). EMBOSS (v6.6) (Rice, Longden, & Bleasby, 2000), Scipio (v1.4.1) (Keller et al., 2008) ( with dependencies in webpage (Scipio, 2025). install BioPerl, Yaml, Blat and the BedOPs function convert2bed ), MAFFT (v 7.490) (Katoh & Standley, 2013) with the option "—auto", and RAxML (v8.2.18) (Stamatakis, 2014) with options set to infer trees from amino acid sequence alignments using an automatic search for the best-fit model of protein evolution and a Gamma rate distribution across sites (PROTGAMMAAUTO), keeping identical sequences (--no-seq-check and –print-identical-sequences) to preserve any gene copies, since HMMER returns non overlapping sequences. These settings can be changed easily, as needed. Instructions for installation of dependencies are provided in the help file in the github.

#### SM4 Input Data Requirements and Test Files

ExTRaCT may be used to search for any conserved single-exon gene or region of interest. ExTRaCT finds genes with known structural motifs and relies on a homolog search program. Four input files are required: the new species genome assembly file(s), a nucleotide homologue sequence FASTA file, a gene protein categorization table and a metadata file. There may be one or many genome assemblies of novel species as targets that have not been annotated or for which coding regions for a certain gene are yet unknown. The FASTA input file with known gene homologues of other species do not need to be in the same taxonomic order or superorder as the target species. The metadata file to initiate the search algorithm requires two columns, the species name and FASTA file path to direct the program. A CSV file with a table of gene labels and corresponding structural motifs is used to filter hits. We show that minor differences in motif definitions greatly impact the count and accuracy of the output.

#### SM5 Pairwise match inspection for all test scenarios

To determine how homologue search results vary depending on input species we conduct three different tests. The first test considers three known species categories, or taxa: bats, Laurasiatheria and primates. A3s may have 1 or 2 catalytic Z-domains which we refer to as single or double domained A3. The second test checks how the algorithm behaves when single, double or combined single and double domain lists are provided as input homologues. We call this last set "both" and includes all A3 for the given species in the taxa. Domain designation varies across mammals. For the bat A3s from Jebb et al. there are 26 genes: 3 double-domain, 19 single-domain (25 total Z-domains) and 4 missing or partial domain A3s. The set of primate A3s consists of three different species (*Homo sapiens*, *Macaca mulatta*, and *Chlorocebus aethiops*), with a total of 9 A3s, of which 4 are double- and 5 are single-domain. The set of Laurasiatheria A3s span 9 species (*Bos taurus*, *Equus caballus*, *Felis catus*, *Lynx lynx*, *Ovis aries*, *Panthera leo bleyenberghi*, *Panthera tigris corbetti*, *Puma concolor*, and *Sus scrofa*), with a total of 25 A3s, with 8 double- and 17 single-domain. The last test uses three different A3 Z-domain amino acid sequence categories defined by Salter (motif set A) (Salter, Bennett, & Smith, 2016), Hayward (motif set B) (Hayward et al., 2018) and Jebb (motif set C) (Jebb et al., 2020).

Salter was the first to classify human cytidine deaminating Z-domains (zinc-dependent deaminase sequence motif (ZDD) in Activation induced deaminase (AID), APOBEC1, APOBEC2 and APOBEC4), and Z1, Z2 and Z3 in A3). Hayward and Jebb define different bat-specific Z-domain structures. In the table we show the specific changes from motif set A to B (Salter to Hayward), then from motif set B to C (Hayward to Jebb). Hayward identified an additional Z2B domain in pteropodids, but also made four changes to Salter motifs, three of which reduce the motif sequence space. Later, Jebb introduced a Z1B domain in bats and made a combination of restrictive and amplifying changes to the sequence space. For our analysis we refer to the input sequence labels and categories as motif set A for Salter, motif set B for Hayward and motif set C for Jebb. Furthermore, the category motifs we defined explicitly restrict the 9th and 10th position for Z1s by excluding the Tryptophan-Phenylalanine (WF) couplet, so sequences with the WF couplet may be categorized as Z2, reflecting the structure of known bat A3 Z2s. For motif set A we treat the 3 canonical Z-domains in Salter as a total of 6 different motifs wherein each domain may have either a Serine or a Threonine before the PC couplet. Rather than providing further novel definitions of Z-domains in bats we chose to compare and use existing definitions which have been verified through expression analysis (Hayward et al., 2018; Jebb et al., 2020).

##### SM5.1. Table. Differences between Motif set categories (A, B, C)

Z domain amino acid sequence motif changes between the three test sets of Salter (A),Hayward (B), and Jebb (C). X marks undefined amino acid in position of motif. Red indicates strict restrictions and green expansions of amino acid sequence space. Arrows indicate increase or decrease of number of unspecified amino acid in motif. Blue with an upward arrow indicates an ambiguous change which expands the number of uncharacterized amino acids but does not include sequences captured by the previous motif. Empty squares are regions of the motif with not change from A to B to C.

| Label | Motif Category | AA Changes | | | | | | | | |
|---|---|---|---|---|---|---|---|---|---|---|
| Z1 | A | | SWS | PC | | F | RIY | | L | |
| | A | | SWT | PC | | F | RIY | | L | |
| | B | | SWX | PC | | F | RIY | | L | |
| | C | ↑ | SWS | PC | | F | RIY | | L | |
| Z1B | C | | SWS | PC | | F | RIY | | L | |
| Z2 | A | | SWS | PC | | F | RLY | | L | |
| | A | | SWT | PC | | F | RLY | | L | |
| | B | | SWS | PC | | F | RLY | | L | |
| | C | ↑ | SWS | PC | | X | RLY | | L | ↓ |
| Z2B | B | | SWS | PC | | F | RIY | | L | |
| | C | | SWS | PC | | F | RIY | | L | |
| Z3 | A | | TWS | PC | | F | RLY | | L | |
| | A | | TWT | PC | | F | RLY | | L | |
| | B | | TWS | PC | | F | RLY | | L | |
| | C | ↓ | TWS | PC | | X | RLX | | L | |

Below are summaries of the 27 test results. We report how the different input tests described, (1) species homologue by clade, (2) gene structure by catalytic arrangement (single domain, double domain and combined single and double domain A3s) and (3) motif filtering definitions (from Salter (Salter, Bennett, & Smith, 2016), Hayward (Hayward et al., 2018) and Jebb (Jebb et al., 2020)), impact results. When relevant, we have listed the amino acid sequences in our three tests below the respective tables.

##### SM5.2. Table.Test 1: Compare input domain files

The following are mismatched sequences found in the first test which compares different input reference sequences by species clade. The reference input A3 sequences domain type is "both" (list of all A3 sequences) and is tested across all classification motifs.

| Sequence key | Species | Found in test result | Missing in test result |
|---|---|---|---|
| Bc1 | *Brachyphylla cavernarum* | Laurasitheria-Both-C | Primate-Both-C |
| Bc1 | *Brachyphylla cavernarum* | Chiroptera-Both-C | Primate-Both-C |

In this case, for domain type "both" and motif set C, Laurasiatheria and chiroptera have identical results, with a total of 496 matching sequences. The 495 sequences from the primate test results are contained in the 496 sequences, but the primate test is missing a sequence in *Brachuphylla cavernarum*, Bc1, which is present in the Laurasiatheria and chiroptera results. Primates are an order of mammals in the superorder Euarchontoglines, and bats are in the order chiroptera, superorder Laurasiatheria. For species of the same superorder (Laurasiatheria) but different order (not chiroptera), the results are the same.

##### SM5.3. Table.Test 2: Compare input domain files

The following are mismatched sequences found in the second test which compare different input reference A3 sequences by domain. The three file types are "both" (all single and double domain A3 sequences), "single" (only list of single domain A3 sequences) and "double" (only list of double domain A3 sequences). The reference input sequences tested are across all species clades and are tested across all classification motifs. List of Li1-4 sequences show a single amino acid difference in sequence outputs.

| Seq key | Species | Found in test result | Missing in test result |
|---|---|---|---|
| Bc1 | *Brachyphylla cavernarum* | Primate-Single-C | Primate-Both-C |
| Ct 1 | *Craseonycteris thonglongyai* | Primate-Both-C | Primate-Single-C |
| Li 1 | *Lonchorhina inusitata* | Primate-Double-A | Primate-Both-A |
| Li 1 | *Lonchorhina inusitata* | Primate-Double-B | Primate-Both-B |
| Li 2 | *Lonchorhina_inusitata* | Primate-Double-A | Primate-Both-A |
| Li 2 | *Lonchorhina_inusitata* | Primate-Double-B | Primate-Both-B |
| Li 3 | *Lonchorhina_inusitata* | Primate-Both-A | Primate-Double-A |
| Li 3 | *Lonchorhina_inusitata* | Primate-Both-B | Primate-Double-B |
| Li 4 | *Lonchorhina_inusitata* | Primate-Both-A | Primate-Double-A |
| Li 4 | *Lonchorhina_inusitata* | Primate-Both-B | Primate-Double-B |

Li Sequences
Li1 – HAEQRFLSWFDDTILSRRAEYQVTWYMSWSPCSECAQEVVAFLEAHENVSLSIFASRLYKCEDEDQQQGLRLLDQAKRTEVAM
Li3 – HAEQRFLSWFDDTILSRRAEYQVTWYMSWSPCSECAQEVVAFLEAHENVSLSIFASRLYKCEDEDQQQGLRLLDQAKRTEVAMM
Li2 – HAEQRFLSWFDGTILSRRAEYQVTWYMSWSPCSECAQEVVAFLEVHENVSLSISASRLYKCEDEEQQQGLRLLDQAKRTEVAM
Li4 – HAEQRFLSWFDGTILSRRAEYQVTWYMSWSPCSECAQEVVAFLEVHENVSLSISASRLYKCEDEEQQQGLRLLDQAKRTEVAMM

Combined domain type input results for "both" include sequences from the same single and double tests, with few exceptions. We would expect that while single and double domain inputs may have different results, if structures vary significantly (for example only double domain have a Z3 but none of the single domain have a Z3), that a combined input file with A3s of both domain types, with all Z types, should result in the same sequences as single- and double- domains separately. We found that, for Laurasiatheria and chiroptera, the third option, a combined list of all A3 genes, captures the same results as when single- and double-domain A3 are analyzed separately. For primate species homologues, one or two sequences were found in single- or double-domain searches that were missing in the corresponding combined domain search.

Test 2 table shows the missing sequences in the domain input test. The *Brachyphylla cavernarum* sequence Bc1 is in the result list from the primate, single-domain, motif set C test, but, as we mentioned, not in the primate combined domain motif set C test. The final hit count of primate single and primate combined domain (motif set C) tests are equal because there is a sequence from *Craseonycteris thonglongyai* (Ct1) in the primate combined domain test result missing from the primate single domain test result. There were two sequences found in the primate double motif set A and B test results missing in the primate combined domain (motif set A and B) tests from the species, *Lonchorhina inusitata* (Li1 and Li2). However, in both motif tests, the primate combined domain results list two other distinct sequences, Li3 which matches the missing Li1, and Li4 which matches the missing Li2, with the addition of a methionine at the end for both. The function nhmmer returns nucleotide sequence ends based on a HMMER profile, so it is likely that in the combined domain case, longer sequences hits were generated to find ORFs. In the case of Bc1, the combined HMMER profile is different enough from the single-domain HMMER profile that it completely missed this sequence, even though the single-domain A3s are a subset of the sequences used to generate the combined-domain HMMER profile. One reason the HMMER profiles of the primates are different enough to miss some results is because fewer sequences were used (9 total) compared to those for the Laurasiatheria (25) and chiroptera (26) HMMER profiles. Both examples (missing Bc1 and varied Li1-4 sequences) show that our algorithm is only weakly sensitive to profile differences, and generally using combined domain inputs will capture the appropriate sequences.

It seems that the number of homologous single- or double-domain genes considered for the pipeline input does not impact the number of output sequences, i.e., more single domains does not necessarily yield more results. For example, there are fewer double-domain than single-domain bat A3s, and in our tests double-domain input consistently yielded fewer results than the single-domain input. Yet the primate A3 homologues we selected had more double-domain than single-domain sequences, albeit a smaller overall count of A3s than either the bat homologue or Laurasiatheria homologue files, and still a similar pattern to those seen with the bat homologues arose in primates, for which the double-domain input yielded fewer results than the single-domain input. Laurasiatheria, in contrast, consistently resulted in more sequences for the double-domain test than the single-domain test, although its count of single- and double-domain A3s resembles the bat homologue input. While HMMER is clearly a robust option for homologue search, it is important to test the different multi-domain inputs separately to capture missing hits, as is the case with the primate species homologue single- and double- domain tests.

##### SM5.4. Table.Test 3 Compare Classification Motif set A, B and C

The following are mismatched sequences found in the third test which compare different classification motifs. The reference input A3 sequence domain file type is "both" (list of all A3 sequences) from the species clade Laurasiatheria. List of Fh1-2 sequences show amino acid differences in sequence outputs.

| Seq key | species | Found in test result | Missing in test result |
|---|---|---|---|
| Li1 | *Lonchorhina inusitata* | Laurasiatheria-Both-C | Laurasiatheria-Both-A |
| Li1 | *Lonchorhina inusitata* | Laurasiatheria-Both-C | Laurasiatheria-Both-B |
| Li2 | *Lonchorhina inusitata* | Laurasiatheria-Both-C | Laurasiatheria-Both-A |
| Li2 | *Lonchorhina inusitata* | Laurasiatheria-Both-C | Laurasiatheria-Both-B |
| Li3 | *Lonchorhina inusitata* | Laurasiatheria-Both-A | Laurasiatheria-Both-C |
| Li3 | *Lonchorhina inusitata* | Laurasiatheria-Both-B | Laurasiatheria-Both-C |
| Li4 | *Lonchorhina inusitata* | Laurasiatheria-Both-A | Laurasiatheria-Both-C |
| Li4 | *Lonchorhina inusitata* | Laurasiatheria-Both-B | Laurasiatheria-Both-C |
| Fh1 | *Furipterus horrens* | Laurasiatheria-Both-B | Laurasiatheria-Both-C |
| Fh2 | *Furipterus horrens* | Laurasiatheria-Both-B | Laurasiatheria-Both-C |

Fh sequences:
Fh1 – HAELCFLKWFEDTILSPYANYAVSWYVSWSPCSSCAEAVVKFLREHKKVKLNIFASRIYYYKYKDKQGLRHLDLAGAQVAMM
Fh2 – HAELCFLQWFEHTILSPYANYDVTWYASWSPCGSCAEAVSTFLREHKKVKLNIFASRIYYYNYKDKQGLRHLVLAGAQVAMM

As mentioned in the main text, motif definition is the most critical input for the algorithm. To avoid underrepresentation of genes while also reducing manual work, the motif filtering and labeling rules are set by the user which provides more flexibility as well as precision. To ensure that the motif set C test hits include all the sequences from the corresponding motif set A and B tests, we limit our pairwise analysis to the results from combined single- and double-domain homologues from the Laurasiatheria test because the same anomalies resulted in the bat and primate tests.

The motif set C results contained all the sequences from the motif set A and B tests, except for 4 sequences; Li3 and Li4 from above (in motif set A and B), and two sequences in *Furipterus horrens,* Fh1 and Fh2 (in motif set A). However, motif set C did pick up Li1 and Li2, the shorter version of Li3 and Li4 missing one of two methionine at the end (**Error! Reference source not found.** Li sequence list). This is because Z2/Z2A defined in motif set C reduced the number of uncharacterized amino acids at the end of the motif. Fh1 and Fh2 correspond to Z2B defined by motif set B but does not match the Z2B motif in set C. This is because of the change from motif set B to C where Z2B has 10 uncharacterized amino acids after the Arginine-Isoleucine-Tyrosine triplet, before the Leucine (RIY-L) instead of 9 (**Error!**

**Reference source not found.**Fh sequence list). Since Fh1 and Fh2 have only 9 amino acids between RIY and L the motif set B test picked them up. Out of all 27 tests, Fh1 and Fh2 are the only sequences missing completely from Laurasiatheria motif set C test. We added these manually to the final gene list table.

#### SM6 Scipio fine tunes nucleotide sequence ends

The Scipio track in our pipeline is optional, as it is meant to correct the start and end locations lost during the ExTRaCT track, however it is a helpful step to gain further insight into the sequence evolution. For example, sequence distances between genes can inform spatial mechanisms in certain evolutionary dynamics. Scipio takes a protein sequence and scans the entire genome assembly to find the corresponding nucleotide sequence. Our algorithm returns the first Scipio match to the final table so if sequences are duplicated in one species, one must look for additional sequence ends in the respective bed files. To run Scipio on the 102 bat genomes and 496 sequences we batched the sequence runs into 14 groups, each containing six or seven species. Running each batch in parallel took approximately 2 hours. To synthesize or characterize a gene, accurate sequence ends are required, hence the addition of this to our overall pipeline.

#### SM7 Validation testing using known bat A3s

To ensure that our algorithm finds true A3s, and identify false negatives, we count the number of known Z-domains from the six bat species in Jebb et al. that were found using ExTRaCT. Motif set C tests returned all the 25 known Z-domains across all clade and domain type tests, except for the double-domain primate A3 homologues test. The two missing Z-domains are both second domains and both Z3s, in the *Rhinolophus ferrumequinum* A3B (Rhifer A3B), and *Rousettus aegyptiacus* A3 (Roueag A3) genes. These two Z3 domains were found in the other Primate motif C tests. This is because none of the double domain primate A3s have a Z3 domain, the only Z3 is in the single domain human A3H, so that the HMMER profiles for double-domain primate A3s will miss homologous similar in structure to Rhifer A3B and Rouaeg A3. Motif set A and B tests had the same results between the two classification techniques, with a high rate of false negatives (22/25) and only 3 matching Z-domains to known bat A3s.

Overall, clade did not impact A3 Z-domain results, and choosing single- or double-domain homologues was only an issue in the primate results. Therefore, we can use ExTRaCT to find accurate A3 Z-domains, with 0 false negatives, if the correct labeling definitions are provided, which is why we proceed with the combined Laurasiatheria A3 motif C results. In the final gene list, we note that among the six bat species (Jebb et al., 2020) an additional eight sequences were found, different from the 25 known bat A3 Z-domains. These may be A3 paralogs or orthologs, depending on when or how the genes diverged. The sequences may be copies from duplication events or speciation events, and they may be non-coding sequences or nonfunctional genes. Species-gene reconciliation can infer whether these genes were separated by speciation (orthologs) or by duplications (paralogs) and further molecular experiments could determine functionality. Importantly, this shows that ExTRaCT finds gene family members that may be missed by conventional annotation methods.

##### SM7.1. Table. Count of known A3 count hits of the six bats

Count of known 6 bat A3s found by ExTRaCT categorized by input species homologue, domain structure and labeling method.

| Hmmer Profile | Domain type | Salter 2016 F1 (HS) | Hayward 2018 F1 (PA) | Jebb 2020 F3 (6B) |
|---|---|---|---|---|
| Bat | Single | 3 | 3 | 25 |
| | Double | 3 | 3 | 25 |
| | Both | 3 | 3 | 25 |
| Primate | Single | 3 | 3 | 25 |
| | Double | 3 | 3 | 23 |
| | Both | 3 | 3 | 25 |
| Laurasiatheria | Single | 3 | 3 | 25 |
| | Double | 3 | 3 | 25 |
| | Both | 3 | 3 | 25 |

#### SM8 Final gene list for phylogenetic analysis

Table by species of all Z domain counts found with ExTRaCT. This list was used

| Species | Z1 | Z2 | Z3 | Total |
|---|---|---|---|---|
| Antrozous pallidus | 3 | 2 | 0 | 5 |
| Artibeus intermedius | 2 | 5 | 0 | 7 |
| Artibeus lituratus | 2 | 2 | 0 | 4 |
| Aselliscus stoliczkanus | 1 | 1 | 1 | 3 |
| Brachyphylla cavernarum | 1 | 3 | 0 | 4 |
| Carollia perspicillata | 1 | 3 | 0 | 4 |
| Centurio senex | 1 | 2 | 0 | 3 |
| Choeroniscus minor | 1 | 3 | 0 | 4 |
| Cistugo seabrae | 2 | 1 | 0 | 3 |
| Corynorhinus mexicanus | 2 | 2 | 0 | 4 |
| Corynorhinus townsendii | 2 | 2 | 0 | 4 |
| Craseonycteris thonglongyai | 1 | 1 | 1 | 3 |
| Cynopterus sphinx | 7 | 2 | 1 | 10 |
| Desmodus rotundus | 1 | 2 | 0 | 3 |
| Diaemus youngii | 2 | 2 | 0 | 4 |
| Diphylla ecaudata | 1 | 2 | 0 | 3 |
| Doryrhina cyclops | 2 | 1 | 1 | 4 |
| Ectophylla alba | 1 | 0 | 0 | 1 |
| Eonycteris spelaea | 3 | 2 | 1 | 6 |
| Eptesicus fuscus | 0 | 2 | 0 | 2 |
| Eptesicus nilssonii | 1 | 3 | 0 | 4 |
| Erophylla bombifrons | 2 | 3 | 0 | 5 |
| Eumops nanus | 0 | 1 | 1 | 2 |
| Furipterus horrens | 1 | 3 | 0 | 4 |
| Glossophaga mutica | 1 | 4 | 0 | 5 |
| Glossophaga soricina | 1 | 4 | 0 | 5 |
| Glyphonycteris daviesi | 1 | 2 | 0 | 3 |
| Hipposideros abae | 0 | 4 | 1 | 5 |
| Hipposideros armiger | 0 | 2 | 1 | 3 |
| Hipposideros caffer | 1 | 4 | 1 | 6 |
| Hipposideros jonesi | 0 | 1 | 1 | 2 |
| Hipposideros larvatus | 0 | 2 | 1 | 3 |
| Hipposideros swinhoii | 1 | 2 | 1 | 4 |
| Hypsignathus monstrosus | 0 | 2 | 1 | 3 |
| Lasiurus ega | 2 | 1 | 0 | 3 |
| Leptonycteris yerbabuenae | 1 | 4 | 0 | 5 |
| Lionycteris spurrelli | 1 | 2 | 0 | 3 |
| Lonchorhina inusitata | 1 | 5 | 0 | 6 |
| Macrophyllum macrophyllum | 1 | 4 | 0 | 5 |
| Macrotus waterhousii | 1 | 4 | 0 | 5 |
| Megaderma spasma | 1 | 2 | 1 | 4 |
| Micronycteris megalotis | 3 | 4 | 0 | 7 |
| Miniopterus australis | 1 | 1 | 0 | 2 |
| Miniopterus natalensis | 1 | 2 | 0 | 3 |
| Miniopterus schreibersii | 1 | 3 | 0 | 4 |
| Molossus alvarezi | 1 | 2 | 1 | 4 |
| Molossus molossus | 1 | 2 | 1 | 4 |
| Molossus nigricans | 1 | 2 | 1 | 4 |
| Mops condylurus | 1 | 2 | 1 | 4 |
| Mormoops megalophylla | 0 | 1 | 0 | 1 |
| Myotis auriculus | 3 | 3 | 0 | 6 |
| Myotis californicus | 0 | 1 | 0 | 1 |
| Myotis daubentonii | 3 | 5 | 0 | 8 |
| Myotis evotis | 2 | 4 | 0 | 6 |
| Myotis lucifugus | 16 | 7 | 0 | 23 |
| Myotis myotis | 1 | 7 | 0 | 8 |
| Myotis mystacinus | 3 | 3 | 0 | 6 |
| Myotis nigricans | 7 | 4 | 0 | 11 |
| Myotis occultus | 12 | 2 | 0 | 14 |
| Myotis pilosus | 4 | 2 | 0 | 6 |
| Myotis velifer | 8 | 4 | 0 | 12 |
| Myotis vivesi | 3 | 2 | 0 | 5 |
| Myotis volans | 1 | 3 | 0 | 4 |
| Myotis yumanensis | 14 | 4 | 0 | 18 |
| Mystacina tuberculata | 2 | 2 | 0 | 4 |

| | | | | |
|---|---|---|---|---|
| Myzopoda aurita | 0 | 1 | 1 | 2 |
| Natalus tumidirostris | 1 | 1 | 1 | 3 |
| Noctilio leporinus | 1 | 2 | 0 | 3 |
| Nyctalus aviator | 4 | 2 | 0 | 6 |
| Nycteris thebaica | 1 | 2 | 2 | 5 |
| Phyllostomus discolor | 2 | 4 | 0 | 6 |
| Phyllostomus hastatus | 3 | 9 | 0 | 12 |
| Pipistrellus kuhlii | 5 | 2 | 0 | 7 |
| Pipistrellus nathusii | 2 | 3 | 0 | 5 |
| Pipistrellus pygmaeus | 3 | 5 | 0 | 8 |
| Platyrrhinus guianensis | 2 | 3 | 0 | 5 |
| Plecotus auritus | 1 | 2 | 0 | 3 |
| Rhinolophus affinis | 1 | 1 | 1 | 3 |
| Rhinolophus ferrumequinum | 2 | 2 | 1 | 5 |
| Rhinolophus foetidus | 1 | 2 | 1 | 4 |
| Rhinolophus hipposideros | 0 | 2 | 1 | 3 |
| Rhinolophus pearsonii | 2 | 1 | 1 | 4 |
| Rhinolophus perniger lanosus | 0 | 2 | 1 | 3 |
| Rhinolophus sinicus | 2 | 1 | 1 | 4 |
| Rhinolophus trifoliatus | 0 | 2 | 1 | 3 |
| Rhinolophus yonghoiseni | 0 | 2 | 1 | 3 |
| Rhinophylla pumilio | 1 | 4 | 0 | 5 |
| Rhinopoma microphyllum | 0 | 1 | 1 | 2 |
| Rhinopoma muscatellum | 0 | 1 | 1 | 2 |
| Rhynchonycteris naso | 2 | 2 | 1 | 5 |

| | | | | |
|---|---|---|---|---|
| Rousettus aegyptiacus | 0 | 2 | 1 | 3 |
| Saccopteryx bilineata | 3 | 1 | 1 | 5 |
| Saccopteryx leptura | 0 | 1 | 1 | 2 |
| Tadarida brasiliensis | 1 | 2 | 1 | 4 |
| Taphozous melanopogon | 1 | 1 | 0 | 2 |
| Thyroptera tricolor | 1 | 6 | 0 | 7 |
| Trachops cirrhosus | 1 | 2 | 0 | 3 |
| Triaenops persicus | 0 | 1 | 1 | 2 |
| Trinycteris nicefori | 1 | 6 | 0 | 7 |
| Uroderma convexum | 2 | 5 | 0 | 7 |
| Vampyressa thyone | 2 | 6 | 0 | 8 |
| Vespertilio murinus | 3 | 3 | 0 | 6 |
| TOTAL | | | | 498 |

**SM9** **MSA for all Z-domains with misclassified sequence (Nt1)**

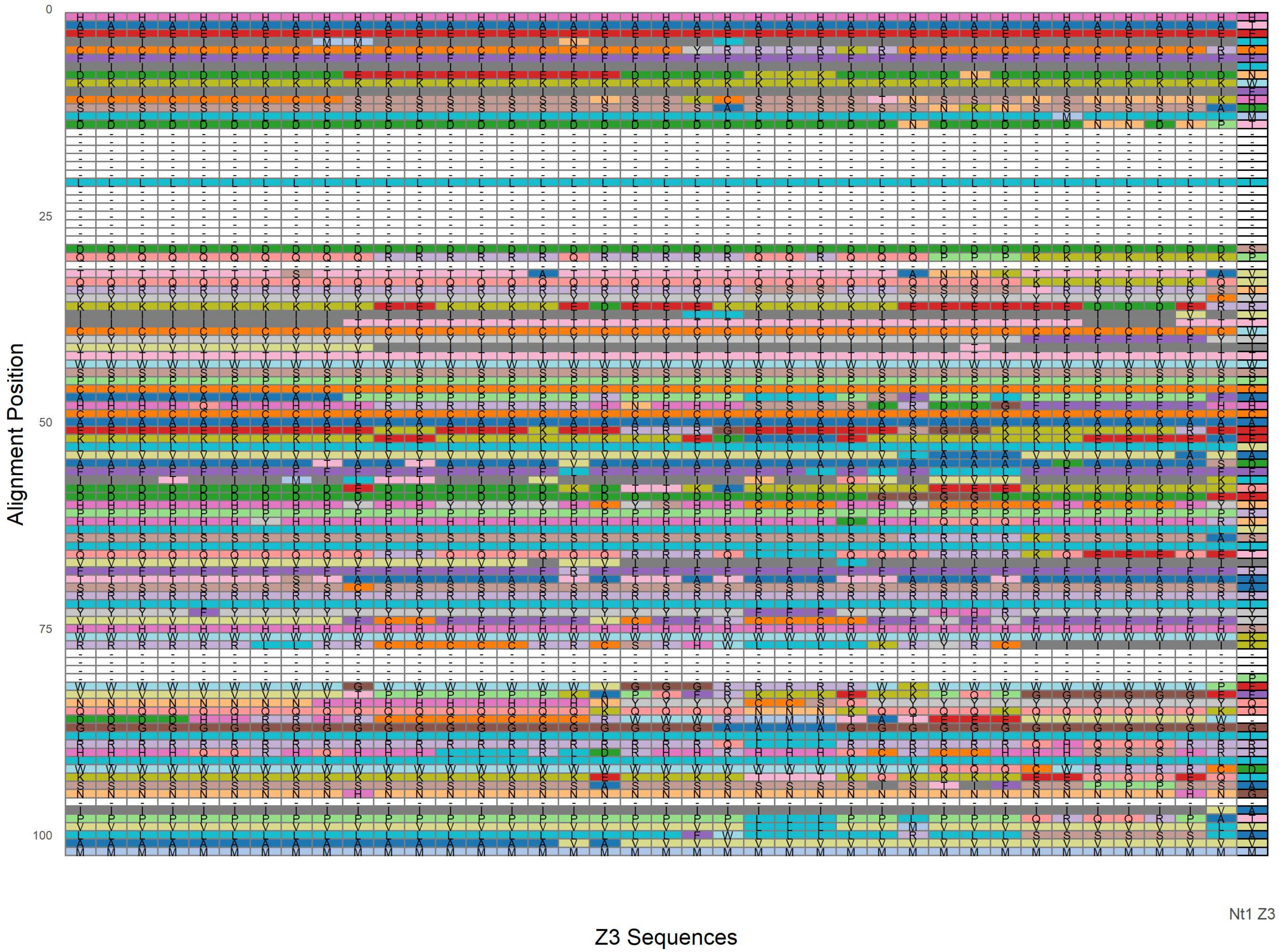

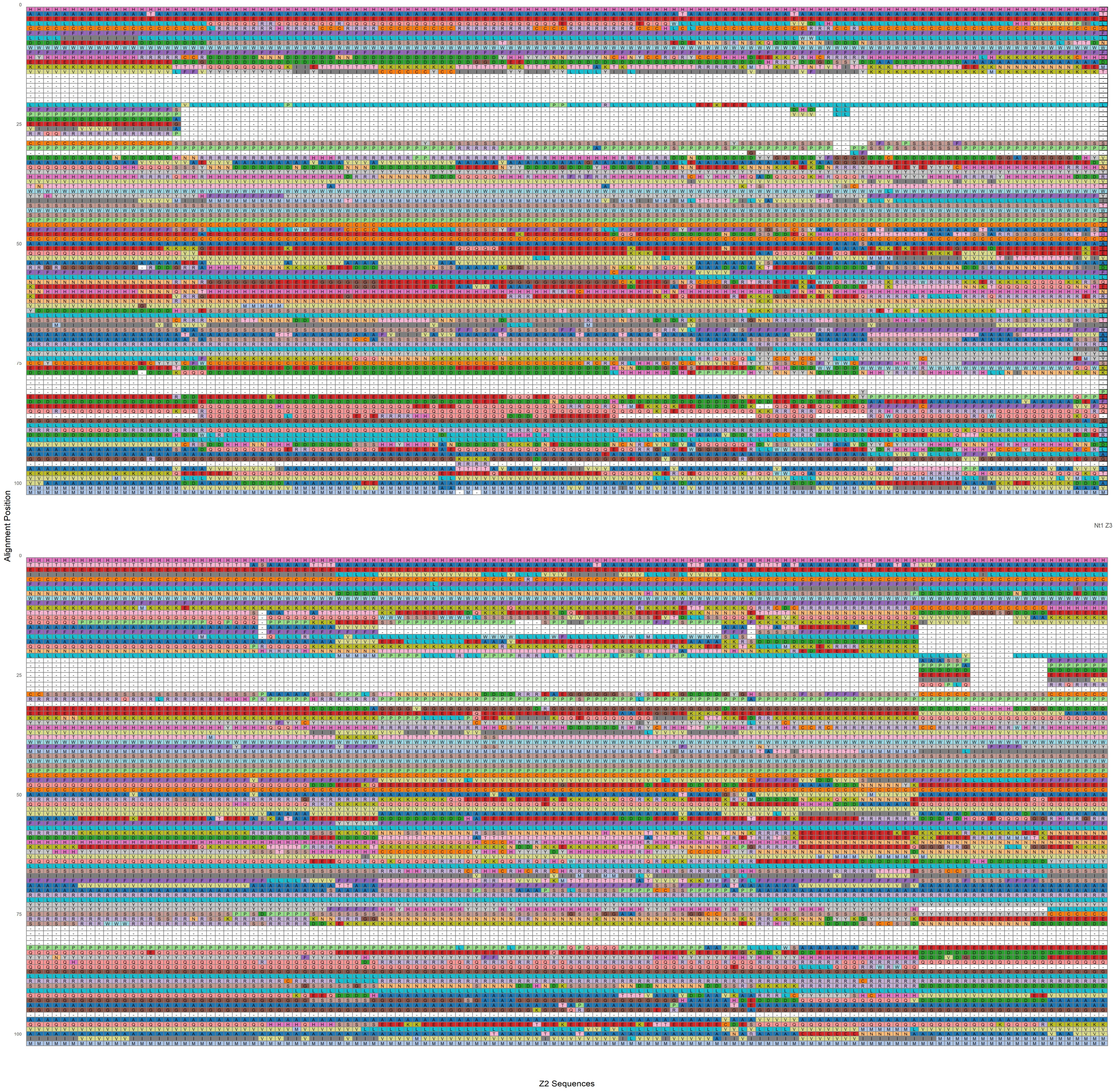
Alignment Position
0
25
50
75
100
Nt1 Z3
0
25
50
75
100
Z2 Sequences

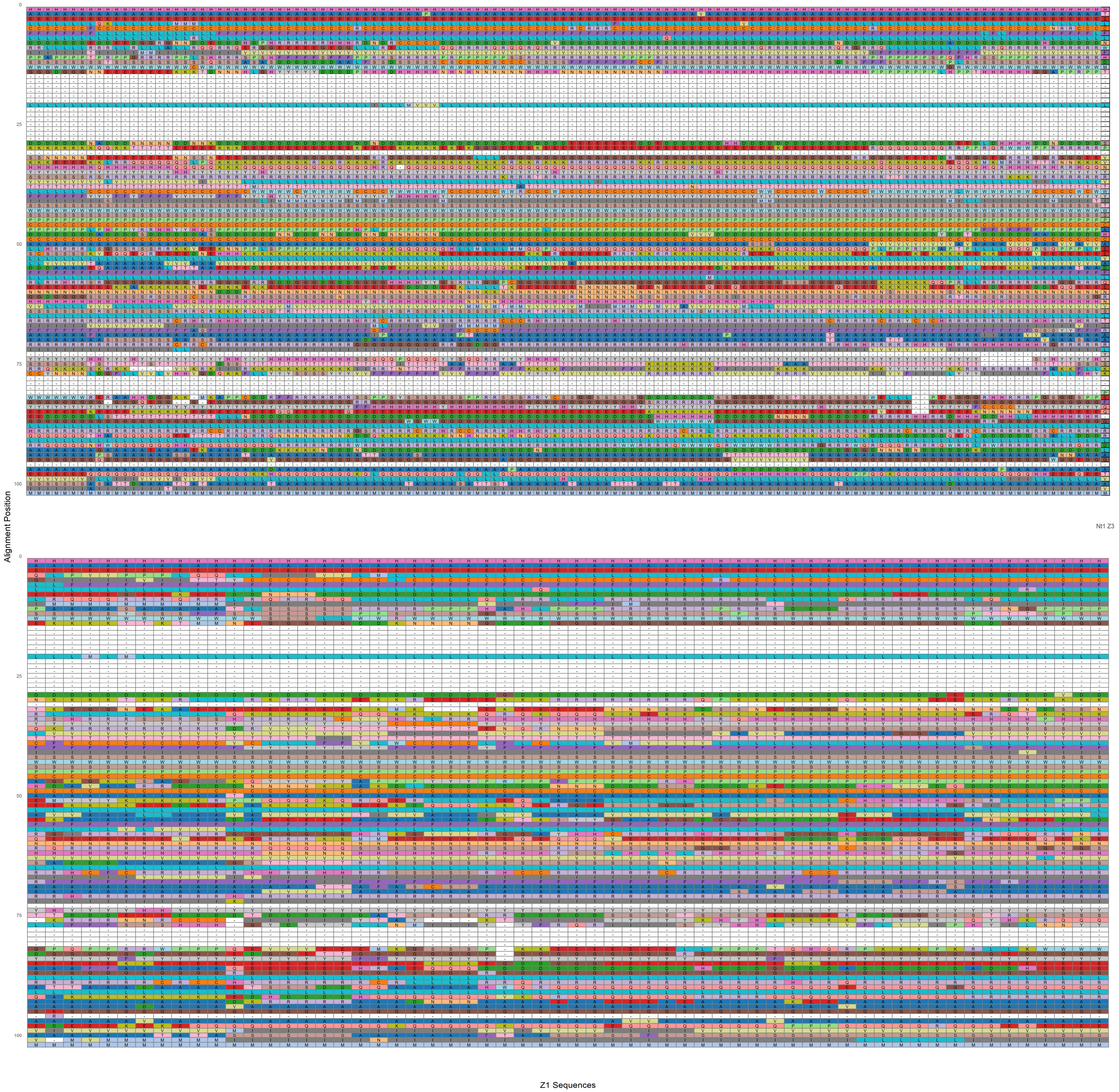
Alignment Position
0
25
50
75
100
Nt1 Z3
Z1 Sequences

**SM10 References**


Banerjee, A., Baker, M. L., Kulcsar, K., Misra, V., Plowright, R., & Mossman, K. (2020). Novel Insights Into Immune Systems of Bats. *Front Immunol*, *11*, 26. https://doi.org/10.3389/fimmu.2020.00026

Bat1K 21 Family Group. (2026). *Bat1K 21 Family Group* Zenodo. https://doi.org/https://doi.org/10.5281/zenodo.19001722

Becker, D. J., Eby, P., Madden, W., Peel, A. J., & Plowright, R. K. (2023). Ecological conditions predict the intensity of Hendra virus excretion over space and time from bat reservoir hosts. *Ecol Lett*, *26*(1), 23-36. https://doi.org/10.1111/ele.14007

Carlson, C. J., Brookson, C. B., Becker, D. J., Cummings, C. A., Gibb, R., Halliday, F. W., Heckley, A. M., Huang, Z. Y. X., Lavelle, T., Robertson, H., Vicente-Santos, A., Weets, C. M., & Poisot, T. (2025). Pathogens and planetary change. *Nature Reviews Biodiversity*, *1*(1), 32-49. https://doi.org/10.1038/s44358-024-00005-w

Cock, P. J., Antao, T., Chang, J. T., Chapman, B. A., Cox, C. J., Dalke, A., Friedberg, I., Hamelryck, T., Kauff, F., Wilczynski, B., & de Hoon, M. J. (2009). Biopython: freely available Python tools for computational molecular biology and bioinformatics. *Bioinformatics*, *25*(11), 1422-1423. https://doi.org/10.1093/bioinformatics/btp163

Dale, R. K., Pedersen, B. S., & Quinlan, A. R. (2011). Pybedtools: a flexible Python library for manipulating genomic datasets and annotations. *Bioinformatics*, *27*(24), 3423-3424. https://doi.org/10.1093/bioinformatics/btr539

Eby, P., Peel, A. J., Hoegh, A., Madden, W., Giles, J. R., Hudson, P. J., & Plowright, R. K. (2023). Pathogen spillover driven by rapid changes in bat ecology. *Nature*, *613*(7943), 340-344. https://doi.org/10.1038/s41586-022-05506-2

Eddy, S. R. (2011). Accelerated Profile HMM Searches. *PLoS Comput Biol*, *7*(10), e1002195. https://doi.org/10.1371/journal.pcbi.1002195

Foundation, P. S. (2021). *Python Programming Language*. In (Version 3.7.10) Python Software Foundation. https://www.python.org/downloads/release/python-3710/

Hayward, J. A., Tachedjian, M., Cui, J., Cheng, A. Z., Johnson, A., Baker, M. L., Harris, R. S., Wang, L. F., & Tachedjian, G. (2018). Differential Evolution of Antiretroviral Restriction Factors in Pteropid Bats as Revealed by APOBEC3 Gene Complexity. *Mol Biol Evol*, *35*(7), 1626-1637. https://doi.org/10.1093/molbev/msy048

Hoyt, B. (2024). *regex*. In (Version 2024.4.16) Python Package Index (PyPI). https://pypi.org/project/regex/

Jebb, D., Huang, Z., Pippel, M., Hughes, G. M., Lavrichenko, K., Devanna, P., Winkler, S., Jermiin, L. S., Skirmuntt, E. C., Katzourakis, A., Burkitt-Gray, L., Ray, D. A., Sullivan, K. A. M., Roscito, J. G., Kirilenko, B. M., Davalos, L. M., Corthals, A. P., Power, M. L., Jones, G., . . . Teeling, E. C. (2020). Six reference-quality genomes reveal evolution of bat adaptations. *Nature*, *583*(7817), 578-584. https://doi.org/10.1038/s41586-020-2486-3

Katoh, K., & Standley, D. M. (2013). MAFFT multiple sequence alignment software version 7: improvements in performance and usability. *Mol Biol Evol*, *30*(4), 772-780. https://doi.org/10.1093/molbev/mst010

Keller, O., Odronitz, F., Stanke, M., Kollmar, M., & Waack, S. (2008). Scipio: using protein sequences to determine the precise exon/intron structures of genes and their orthologs in closely related species. *BMC Bioinformatics*, *9*, 278. https://doi.org/10.1186/1471-2105-9-278

NCBI Resource Coordinators. (2016). Database resources of the National Center for Biotechnology Information. *Nucleic Acids Res*, *44*(D1), D7-19. https://doi.org/10.1093/nar/gkv1290

O'Shea, T. J., Cryan, P. M., Cunningham, A. A., Fooks, A. R., Hayman, D. T., Luis, A. D., Peel, A. J., Plowright, R. K., & Wood, J. L. (2014). Bat flight and zoonotic viruses. *Emerg Infect Dis*, *20*(5), 741-745. https://doi.org/10.3201/eid2005.130539

Perez, G., Barber, G. P., Benet-Pages, A., Casper, J., Clawson, H., Diekhans, M., Fischer, C., Gonzalez, J. N., Hinrichs, A. S., Lee, C. M., Nassar, L. R., Raney, B. J., Speir, M. L., van Baren, M. J., Vaske, C. J., Haussler, D., Kent, W. J., & Haeussler, M. (2025). The UCSC Genome Browser database: 2025 update. *Nucleic Acids Res*, *53*(D1), D1243-D1249. https://doi.org/10.1093/nar/gkae974

Reback, J., jbrockmendel, McKinney, W., Van den Bossche, J., Augspurger, T., Cloud, P., Hawkins, S., Roeschke, M., gfyoung, Sinhrks, Klein, A., Petersen, T., Hoefler, P., Tratner, J., She, C., Ayd, W., Naveh, S., Garcia, M., Darbyshire, J., & Seabold, S. (2021). *pandas-dev/pandas: Pandas 1.3.5 (v1.3.5)*. In

Rice, P., Longden, I., & Bleasby, A. (2000). EMBOSS: the European Molecular Biology Open Software Suite. *Trends Genet*, *16*(6), 276-277. https://doi.org/10.1016/s0168-9525(00)02024-2

Salter, J. D., Bennett, R. P., & Smith, H. C. (2016). The APOBEC Protein Family: United by Structure, Divergent in Function. *Trends Biochem Sci*, *41*(7), 578-594. https://doi.org/10.1016/j.tibs.2016.05.001

Scipio, T. (2025). *Scipio eukaryotic gene identification*. In https://www.webscipio.org/help/scipio#requirements

Stamatakis, A. (2014). RAxML version 8: a tool for phylogenetic analysis and post-analysis of large phylogenies. *Bioinformatics*, *30*(9), 1312-1313. https://doi.org/10.1093/bioinformatics/btu033

Stanke, M., & Morgenstern, B. (2005). AUGUSTUS: a web server for gene prediction in eukaryotes that allows user-defined constraints. *Nucleic Acids Res*, *33*(Web Server issue), W465-467. https://doi.org/10.1093/nar/gki458

UCSC. (2025). *Senckenberg Genome Browser*. https://genome.senckenberg.de/

Zhang, G., Cowled, C., Shi, Z., Huang, Z., Bishop-Lilly, K. A., Fang, X., Wynne, J. W., Xiong, Z., Baker, M. L., Zhao, W., Tachedjian, M., Zhu, Y., Zhou, P., Jiang, X., Ng, J., Yang, L., Wu, L., Xiao, J., Feng, Y., . . . Wang, J. (2013). Comparative analysis of bat genomes provides insight into the evolution of flight and immunity. *Science*, *339*(6118), 456-460. https://doi.org/10.1126/science.1230835
